# Lipidomic analysis of human TANGO2-deficient cells suggests a lipid imbalance as a cause of TANGO2 deficiency disease

**DOI:** 10.1101/2024.02.29.581305

**Authors:** Mahsa Mehranfar, Paria Asadi, Rozmehr Shokohi, Miroslav P. Milev, Chiara Gamberi, Michael Sacher

## Abstract

TANGO2 deficiency disease (TDD) is a multisystem disease caused by variants in the *TANGO2* gene. Symptoms include neurodevelopmental delays, seizures and potentially lethal metabolic crises and cardiac arrhythmias. While the function of TANGO2 remains elusive, vitamin B5/pantothenic acid supplementation has been shown to alleviate symptoms in a fruit fly model and has also been used to treat individuals suffering from TDD with success. Since vitamin B5 is the precursor to the lipid activator coenzyme A (CoA), we hypothesized that TANGO2-deficient cells would display changes in the lipid profile compared to control and that these changes would be rescued by vitamin B5 supplementation. In addition, the specific changes seen might point to a pathway in which TANGO2 functions. Indeed, we found profound changes in the lipid profile of human TANGO2-deficient cells as well as an increased pool of free fatty acids in both human cells devoid of TANGO2 and *Drosophila* harboring a previously described TANGO2 loss of function allele. All these changes were reversed upon vitamin B5 supplementation. Pathway analysis showed significant increases in triglyceride as well as in lysophospholipid levels as the top enriched pathways in the absence of TANGO2. Consistent with a defect in triglyceride metabolism, we found changes in lipid droplet numbers and sizes in the absence of TANGO2 compared to control. Our data will allow for comparison between other model systems of TDD and the homing in on critical lipid imbalances that lead to the disease state.

## INTRODUCTION

*TANGO2* was first discovered as a gene whose product was important for Golgi transport and organization [1]. However, the protein was found to be cytosolic with a portion bound to mitochondria, suggesting its role in Golgi transport may be indirect [2,3]. Since the first reports of individuals with mutations in TANGO2 leading to TANGO2 deficiency disease (TDD) [4,5], nearly 100 individuals with a number of putatively dysfunctional variants have been described [6]. Though originally thought to be a mitochondrial disease, mito-cocktail treatment has proven to be ineffective [7], and affected individuals were thought to have defects in some aspect of fatty acid metabolism [4]. Among the most common phenotypes in TDD are intellectual deficit, neurodevelopmental delay, seizures, and illness- or starvation-induced metabolic crises that lead to potentially lethal cardiac arrhythmias [7]. Thus, alleviating the possibility of a metabolic crisis is crucial in reducing mortality and morbidity.

We have established a TDD model in *Drosophila* fruit flies and have shown this model recapitulates many of the phenotypes seen in TDD-affected individuals [8]. Importantly, these phenotypes are alleviated by treatment with vitamin B5/pantothenic acid. This supplementation also restored normal Golgi transport in human fibroblasts devoid of this protein [8]. Since the only known function of vitamin B5 in humans is as a precursor to the production of coenzyme A (CoA), a lipid “activator” involved in many aspects of lipid metabolism, we speculated that the lipid profile in both human and fruit fly TANGO2 deficient cells would be affected. We further hypothesized that this imbalance would be corrected by vitamin B5 supplementation. Here, we put these notions to the test and indeed find significant changes in the lipid profile of TANGO2-deficient cells. These changes were largely reversed by vitamin B5 treatment, and pathway analysis suggests that triglyceride metabolism and production of phospholipids by lysophospholipids are the most profoundly affected pathways. Consistent with a triglyceride effect, we find that a portion of human TANGO2 associates transiently with lipid droplets during glucose deprivation. We discuss our data in the context of lipid analyses of other model systems recently reported.

## MATERIALS AND METHODS

**(*Note that detailed methods can be found in the supplementary information file that accompanies this study*)**

### Cell culture and fly husbandry

Human fibroblasts and HepG2 cells were grown in Dulbecco’s modified Eagle medium (DMEM) supplemented with 10% fetal bovine serum (FBS). If cultures were treated with vitamin B5, this vitamin was used at 2 mM for 4 days prior to harvesting for lipid analysis. Flies were maintained at room temperature on Bloomington formulation food either with or without 4 mM vitamin B5. Free fatty acid analysis was performed on 10-12 day old flies.

### Lipid analysis

Cells or flies were harvested and washed in PBS. Mass spectrometry-based lipid (human) and free fatty acid analyses (human, *Drosophila*) was performed by Lipotype GmbH (Dresden, Germany) using a chloroform/methanol extraction procedure. Detailed methods can be found in the supplemental information section.

### Lipid droplet staining and microscopy

HepG2 cells were transfected with TANGO2-RFP and then treated with 200 uM oleic acid for 6 hours to generate lipid droplets. Oleic acid was removed, and the medium replaced with either DMEM/10% FBS or glucose-free DMEM/10% FBS. The cells were fixed in 4% paraformaldehyde at various time points after washout and fixed. Lipid droplets were stained with BODIPY 495/503. Images were acquired using an Olympus FIV10i confocal laser-scanning microscope with a 60X objective lens (N.A. 1.35). Images were analyzed using Image J with the JACoP plugin.

### Western analysis of TANGO2 levels during starvation

Cells were grown in glucose-free DMEM/10% FBS with 10 mM galactose for up to 15 days. The cells were harvested in lysis buffer (10 mM Tris-HCl pH 8.0, 1 mM EDTA, 1% Triton X-100, 0.1% sodium deoxycholate, 0.1% SDS with protease inhibitors and 10 mM PMSF). Total protein (20 μg) was fractionated by SDS-PAGE, transferred to PVDF membranes and probed with anti-TANGO2 (homemade) and anti-tubulin (Sigma). After visualization by chemiluminescence, bands were quantified using ImageJ and TANGO2 expression was normalized to tubulin. The experiment was repeated 3 times and the average ± SEM was plotted.

### Pathway analysis

Lipid data generated above was subjected to pathway analysis using Lipid Network Explorer (Linex^2^) [9] based on the Rhea and Reactome databases.

## RESULTS AND DISCUSSION

Human fibroblasts devoid of TANGO2 and control fibroblasts were subjected to lipid profiling. As seen in Figure 1A, significant changes were noted between untreated control and TANGO2-deficient fibroblasts (see supplemental file 1 for the raw data of the lipid profiles and class-specific changes). Chief among these changes was an increase in triacylglyceride (TAG) levels (Figure 1B). This change was exacerbated when cells were deprived of glucose (Figures 1C,D). Upon treatment of the cells with 2 mM vitamin B5 for 4 days prior to analysis, the levels of accumulated TAGs significantly decreased. Other significant lipid accumulations seen in TANGO2-deficient human cells were also largely rescued following vitamin B5 treatment (Figure 1E,F). We further noted an increase in free fatty acids (supplemental Figure S1; full raw data in supplemental file 2), consistent with an increase in lysophospholipids seen in the absence of TANGO2 (Figure 1C,D). The increase in free fatty acids was also seen in a Drosophila model of TDD (supplemental Figure S1), though a full lipid analysis has yet to be undertaken.

**Figure 1:**
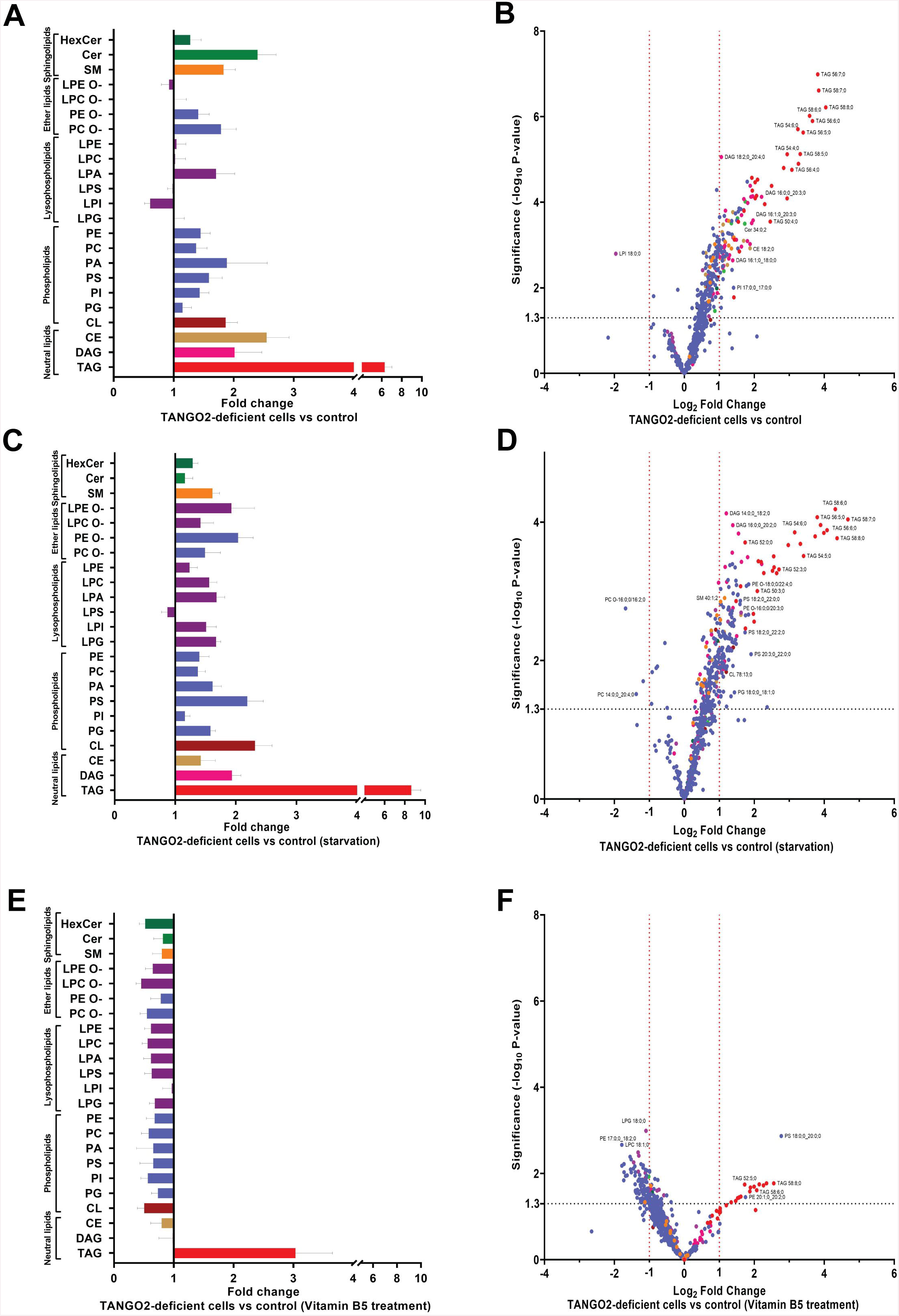
Lipid changes in human fibroblasts are rescued by vitamin B5 supplementation. Control and TANGO2 deficient fibroblasts were subjected to lipidomics analysis as described in the methods section. Data was plotted as the ratio of TANGO2 deficient cells to control as lipid classes (A, C, E) and in volcano plots (B, D, F). Cells were either untreated (A, B), starved of glucose (C, D) or treated with vitamin B5 (E, F). Colours between panels A, C and E are colour-matched to panels B, D and F. The horizontal dashed line in the volcano plots represents p=0.05.

We then subjected the human lipid profile to pathway analysis using Lipid Network Explorer 2 (Linex^2^) [9]. This analysis revealed many potential lipid processes that were affected by the absence of TANGO2 (see supplemental Figure S2A). Pathway enrichment was performed to highlight the most significantly affected pathways. As seen in Figure 2A, pathways affecting TAG synthesis were the most highly enriched. Also significantly enriched were pathways leading to the production of lysophospholipids such as lysophosphatidylinositol and lysophosphatidylcholine (Figure 2B). When we examined the accumulated TAGs more carefully, we noted that the increase was more skewed for the polyunsaturated TAGs of ≥54 total carbons (Figure 2C). Upon treatment of the cells with vitamin B5, many of these pathway changes were rescued (supplemental Figure S2B). These results are consistent with TANGO2 functioning in some aspect of lipid metabolism in human fibroblasts, perhaps TAG metabolism.

**Figure 2:**
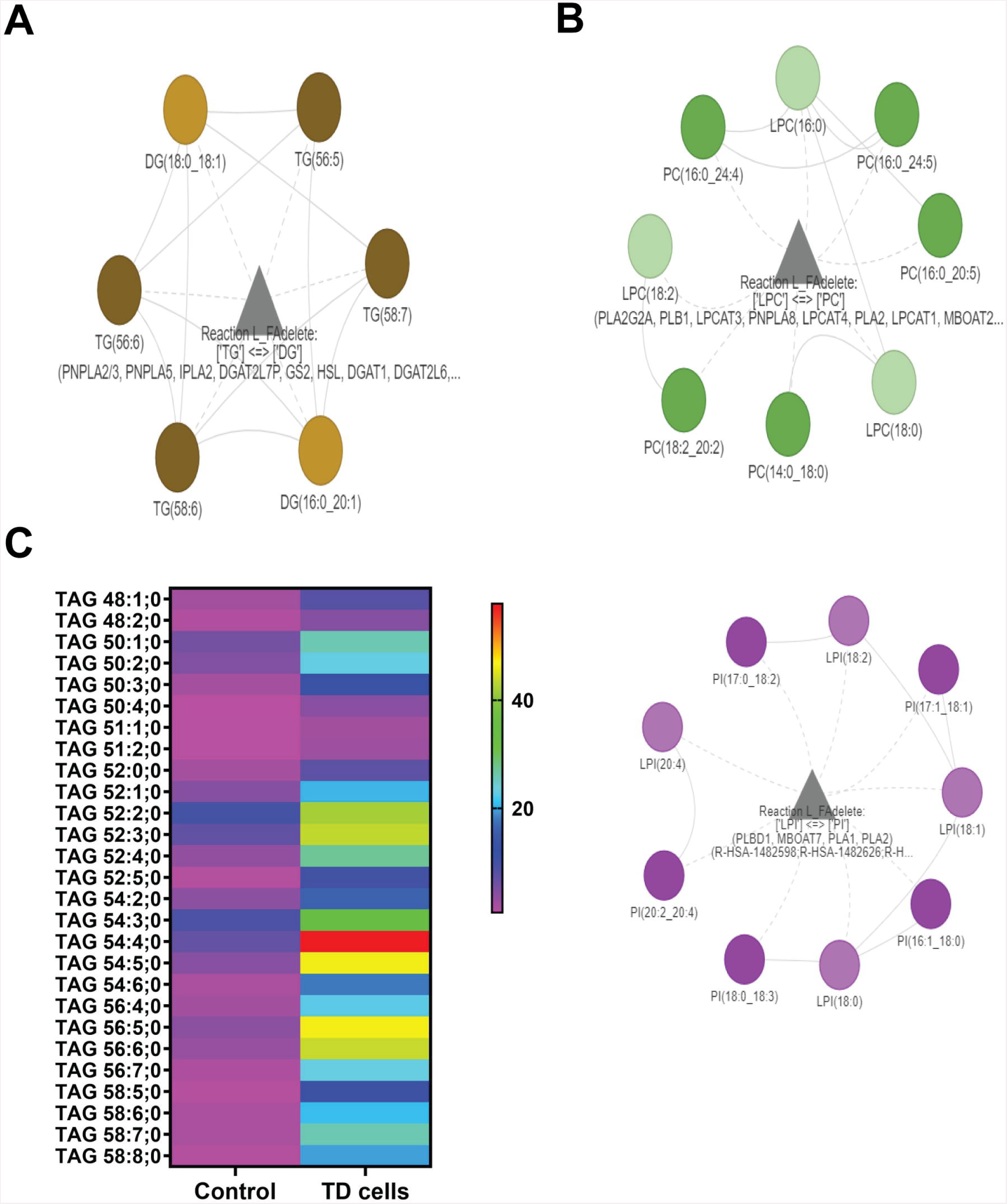
Pathway analysis of human lipidomics data suggests an effect on triacylglycerides. The lipidomics data for untreated (A) or vitamin B5 supplemented cells (B) was subjected to pathway analysis as described in the methods section. The full analysis is shown in Figure S2. The most significantly enriched pathways are shown in panels A and B, affecting triacylglycerides (A) and lysophospholipids (B). (C) A heat map of triacylglycerides accumulating TANGO2 deficient (TD) cells is shown.

Given the fact that starvation exacerbates the lipid imbalance seen in TANGO2-deficient cells, we examined whether the human cells have a stronger need for TANGO2 during glucose starvation when cells have an increased need to metabolize lipids. As shown in Figure 3A and B, we found a time-dependent increase in TANGO2 levels during glucose starvation in human fibroblasts. This effect has also been observed in several different cell types (Figure S3) and is not specific to fibroblasts. These results are consistent with TANGO2 having a function in lipid metabolism that is required when cellular lipid metabolism needs are increased.

**Figure 3:**
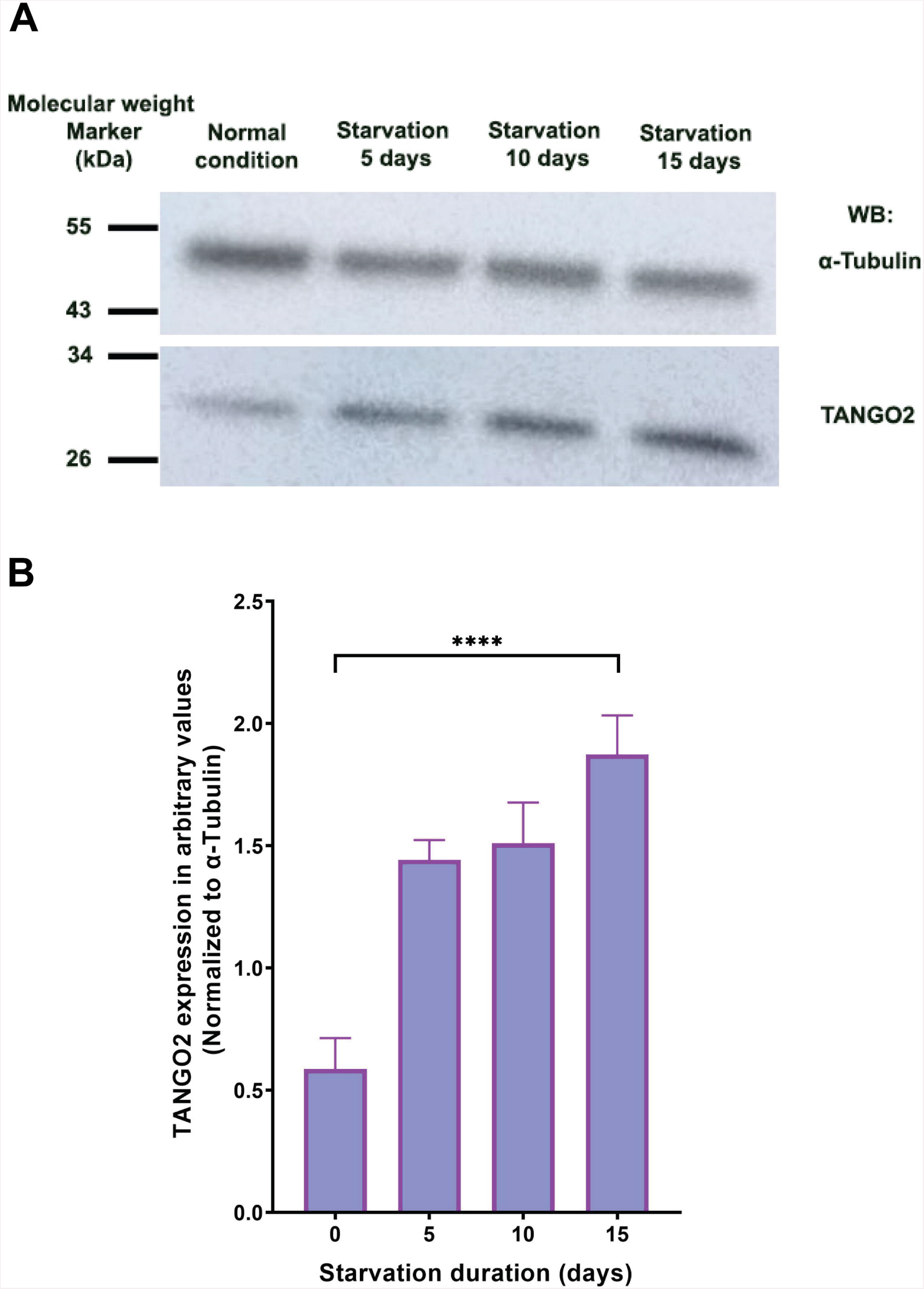
TANGO2 levels increase during glucose deprivation in fibroblasts. Control fibroblasts were either unstarved (normal conditions) or deprived of glucose in the presence of galactose for the times indicated. TANGO2 and tubulin (loading control) were visualized by western analysis (A) and quantified (B). The experiment was repeated 3 times and a representative blot is shown. For statistical significance, **** indicates P-value<0.0001.

Having seen a striking increase in TAG levels in the absence of functional TANGO2, and since TAGs are stored in lipid droplets (LDs), we examined whether TANGO2 associates with LDs. We speculated that TANGO2 may associate with LDs during conditions when LDs are metabolized. Therefore, we generated LDs by treating HepG2 cells with oleic acid, and then assessed the association of fluorescently-tagged TANGO2 (TANGO2-RFP) with the LDs during oleic acid washout, conditions where LDs will be consumed by the cell. The LDs were visualized using the fluorescent marker BODIPY. As shown in Figure 4A and quantified in Figure 4B, we noted a small but significant association of TANGO2-RFP with LDs within 0.5 hours of oleic acid washout, peaking after 1 hour of washout. If TANGO2 is involved in LD metabolism, then we expect to see changes in LDs in the absence of TANGO2. Using human fibroblast cells devoid of TANGO2 protein [3], we measured LD numbers before and after stimulation with oleic acid.

**Figure 4:**
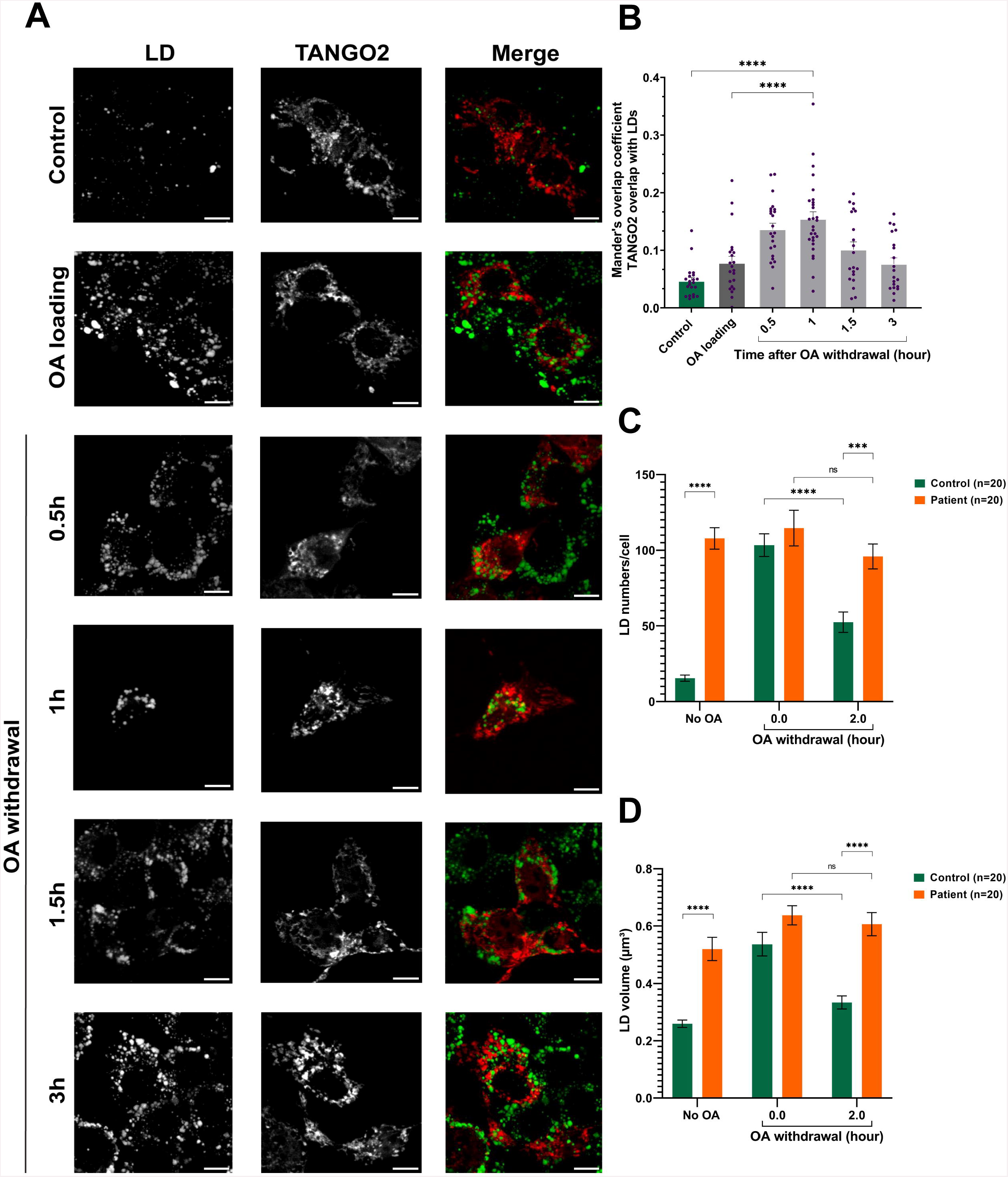
TANGO2 associates transiently with lipid droplets in HepG2 cells and affects lipid droplet numbers and sizes in human fibroblasts. (A) HepG2 cells were transfected with TANGO2-RFP and untreated or treated with oleic acid. Following washout of oleic acid, cells were either fixed immediately or maintained in glucose-deprived medium for the time points indicated. TANGO2 and lipid droplets were then stained with BODIPY and visualized by confocal microscopy. Lipid droplets (green) that were closely associated with TANGO2-RFP (red) were determined using ImageJ. The values were quantified in panel B. The scale bar represents 10 μm. N values are 20-30 cells analyzed over 2 biological replicates and 2 technical replicates. (C, D) Fibroblasts devoid of TANGO2 or control fibroblasts were untreated or treated with oleic acid for 24 hours. Oleic acid was then washed out and cells were immediately fixed or fixed after 2 hours washout and stained with BODIPY. Images were taken and processed for lipid droplet number (C) and for lipid droplet volume (D) using ImageJ. For statistical significance, **** indicates P-value<0.0001, *** indicates P-value<0.001. NS indicates not significant.

As shown in Figure 4C and Figure S4, TANGO2-deficient fibroblasts had more LDs compared to control prior to oleic acid treatment. Oleic acid treatment increased the number of LDs in control, and within 2 hours of oleic acid washout, the LD number dropped significantly in control but not in TANGO2 deficient cells. We also measured LD volume and found that it increased in control cells after oleic acid treatment but was significantly decreased following washout in controls but not in TANGO2 deficient cells (Figure 4D and Figure S4). These results suggest that TANGO2 affects LD biogenesis and degradation.

Recent studies have examined the lipid profile in other TDD models including zebrafish and HepG2 (hepatocellular carcinoma) knockdown cells [10,11]. In the zebrafish model, TAGs were decreased as opposed to the ∽6-fold increase we noted in human fibroblasts. Furthermore, phosphatidylcholine (PC) and phosphatidylethanolamine (PE) were decreased in the zebrafish model. Surprisingly, the precursors lysopho-PC and -PE (LPC and LPE), were also found to be decreased. These observations contrast to our data in human fibroblasts showing increased LPC and free fatty acid levels. Similar to our study, in the HepG2 TANGO2 knockdown model [11], a significant increase in lysophosphatidic acid (LPA) was noted with a concomitant decrease in PA. Similarly, most other phospholipids showed increases in the lysophospholipids and a decrease in the corresponding phospholipid. The authors also noted a decrease in cardiolipin, a mitochondrial-specific lipid, possibly providing an explanation for mitochondrial defects that have been periodically reported in TANGO2-deficient cells. We did not observe a decrease in cardiolipin in our human fibroblast model system. Furthermore, the authors did not note the same striking increase in TAG levels as we detected in the TANGO2-deficient cells.

Comparison of the lipidomics data from our study and previously published studies did not reveal a lipid “signature” of species consistently affected across the various model systems. While these studies collectively reinforce the notion that TANGO2 dysfunction affects lipid balance in cells, which is likely causative for TDD, how this imbalance occurs and why it results in the disease state remains unknown. We can speculate on several possibilities. First, TANGO2 may have a broad function in lipid metabolism that varies between the cell type and model system used. Perhaps in human fibroblasts the protein is involved in TAG metabolism, though it is unclear why this would not be conserved in fish or hepatocellular carcinoma cells (HepG2) considering TANGO2 is a highly evolutionarily conserved protein. The increase in lysophospholipids seen in both human models supports a role for TANGO2 in biogenesis of this lipid. Second, since lipid and physiological defects are rescued by vitamin B5 administration [8,12-14], and since vitamin B5 is responsible for producing coenzyme A (CoA) in a highly conserved five-step pathway, TANGO2 may be required for some aspect of CoA metabolism. CoA levels are tightly regulated in cells [15], with constant synthesis and degradation resulting in the proper balance. Although defects in CoA synthesis result in neurodegeneration with brain iron accumulation (NBIA; OMIM 617917), this characteristic iron accumulation in NBIA has not been reported in TDD-affected individuals. Consistent with an imbalance in CoA as a possible cause of TDD, it is noteworthy that defects in the mitochondrial CoA transporter SLC25A42 (OMIM 618416) have phenotypes similar to those of TDD-affected individuals including stress- or illness-induced metabolic crises, seizures, neurologic regression, hypotonia and impaired movement [16,17]. Clearly, further work is necessary to determine the precise function(s) of TANGO2, and our lipidomic data from human cells will serve as a necessary dataset in these endeavors.

## ACKNOWLEDGEMENTS

We are grateful to Dr. Dajana Vuckovic for initial consultations on the lipidomics work. We are also grateful to the CBAMS and CMCI facilities at Concordia University for use of the mass spectrometry and microscopy equipment, respectively. This work was funded by the TANGO2 Research Foundation (to MS and CG), the Canadian Institutes of Health Research and the Natural Sciences and Engineering Research Council (to MS).

## AUTHOR CONTRIBUTIONS

MM prepared cells for lipidomics analysis, interpreted the data, performed pathway analysis and all microscopy, and edited the manuscript. PA performed the fly work for fatty acid analysis and edited the manuscript. MPM performed the analysis of TANGO2 levels in HeLa and HEK293 cells. RS prepared human fibroblasts for the data in Figures 4 and S3, and edited the manuscript. CG interpreted the data, oversaw the fly work and edited the manuscript. MS wrote and edited the manuscript, interpreted the data, conceptualized and oversaw the entire study.

