## Supplementary figures and images for "Lipidomic analysis of human TANGO2-deficient cells suggests a lipid imbalance as a cause of TANGO2 deficiency disease"

### Figure S1

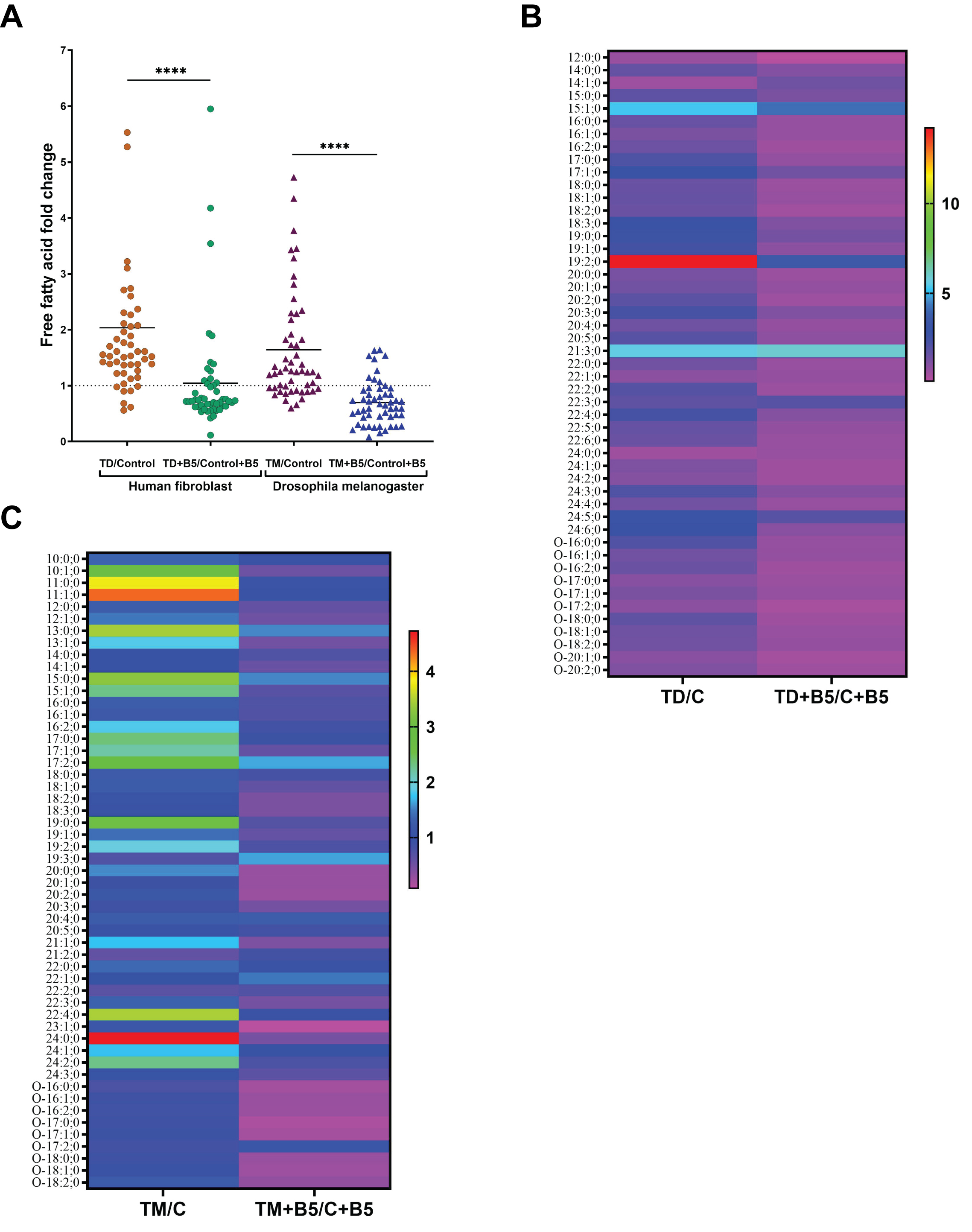

### Figure S2

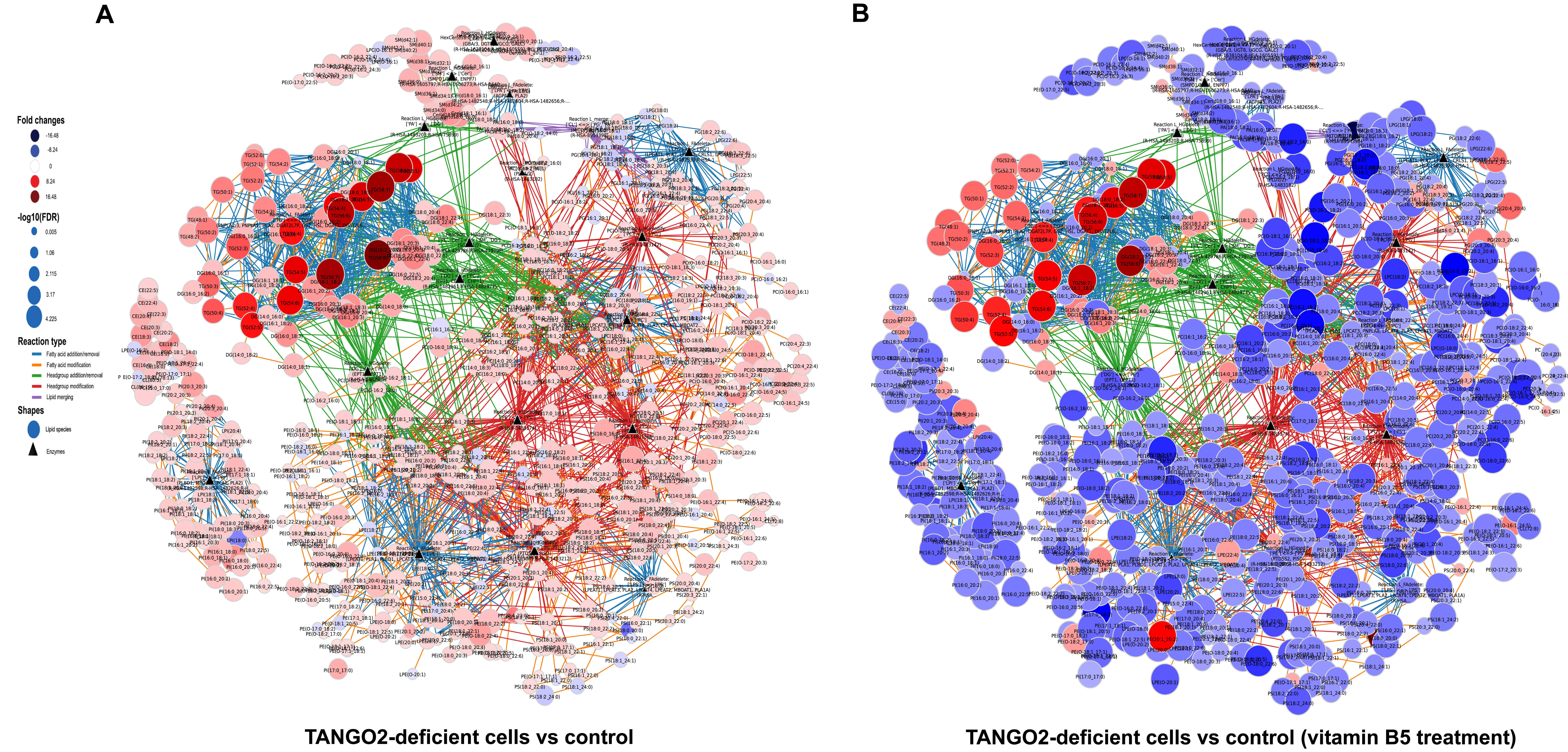

### Figure S3

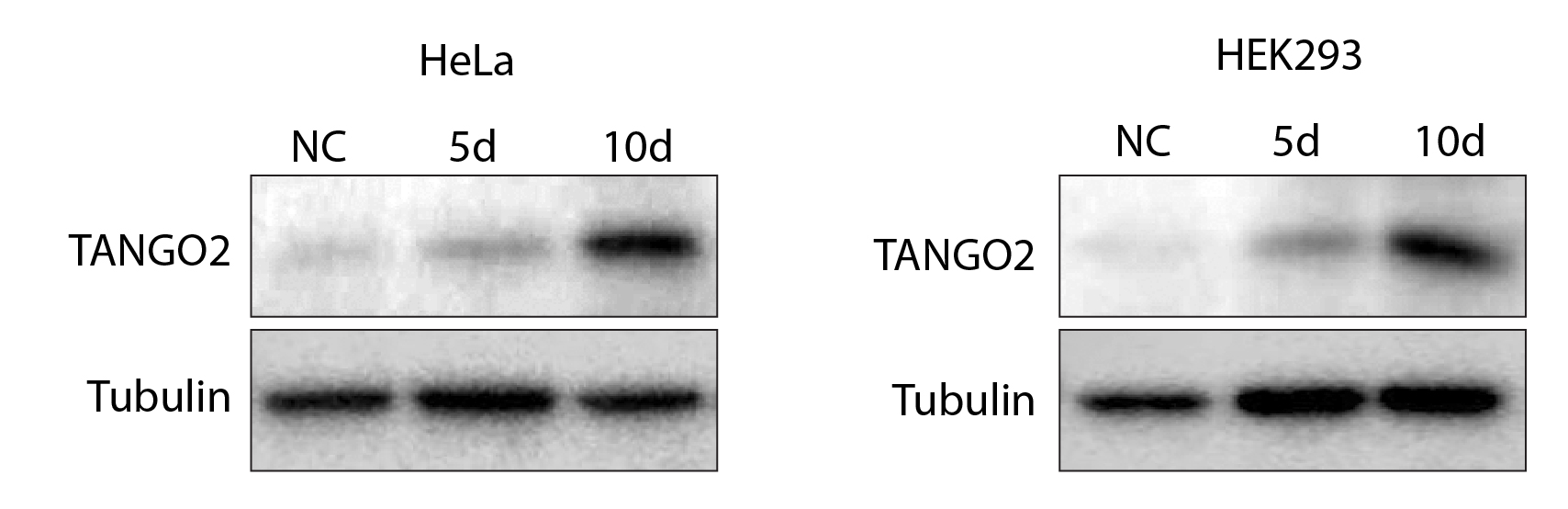

### Figure S4

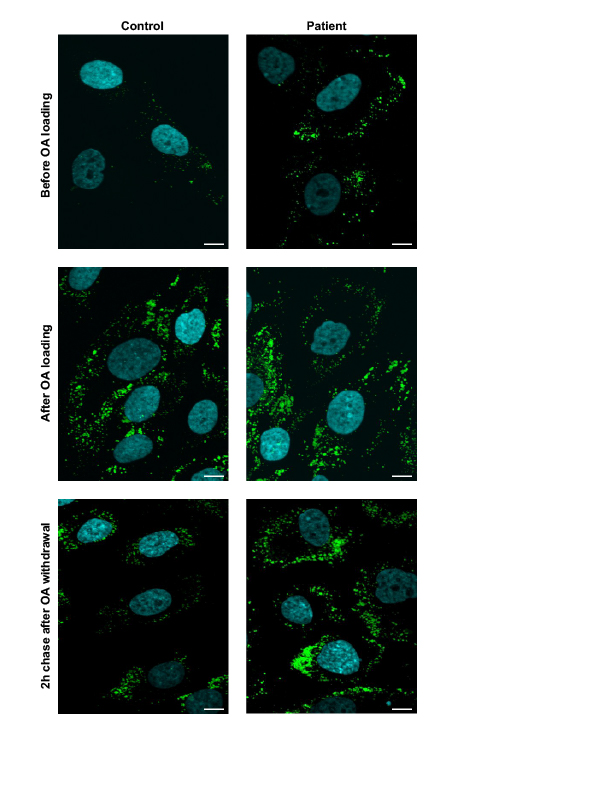
